# Malaria parasite population genomics during an elimination program in Eastern Myanmar

**DOI:** 10.1101/2025.05.21.655408

**Authors:** Xue Li, Grace A. Arya, Aung Myint Thu, Jordi Landier, Daniel M. Parker, Gilles Delmas, Ann Reyes, Khin Maung Lwin, Kanlaya Sriprawat, François Nosten, Timothy J.C. Anderson

## Abstract

**Background:** We investigated parasite population genomics during an intensive malaria elimination program in Kayin State (Myanmar), in which malaria posts were used for rapid detection and treatment of malaria cases, while mass drug administration (MDA) was used in villages with high submicroscopic reservoirs.

**Methods:** We collected 5014 dried blood spots from *Plasmodium falciparum* infected patients from 413 malaria posts, over 58-months (November 2015 - August 2020), and sequenced 2270 parasite genomes, each with geographic references (latitude and longitude). We used identity-by-descent (IBD) relationships to examine how control efforts impact parasite population structure.

**Findings:** Parasites were genetically depauperate: 1726 single-genotype infections comprised 166 unique genomes (≥90% IBD), while nine families (≥45% IBD) accounted for 62·5% of parasites sampled. We observed localized, temporally stable transmission of unique parasite genotypes, identifying transmission chains. Parasite relatedness was positively correlated up to approximately 20 km revealing the scale of parasite subpopulations. *Kelch13* diversity was stable from 2016-2019, but only one predominant clonal genotype (*kelch13*-R561H) remained in 2020. MDA resulted in parasite founder effects, providing genomic evidence for the efficacy of this malaria control tool.

**Interpretation:** Our genomic data show that parasite population size decreased over the study period, and we observed regional distribution of parasite genotypes, which can define operational units for parasite control. One parasite genotype (*kelch13*-R561H) from the north rose to high frequency in 2020, because transmission was halted elsewhere in the control area. Future surveillance will reveal whether this genotype spreads to neighboring regions. Genetic drift may have a stronger impact on parasite population structure than selection in low-transmission elimination settings.

**Funding:** National Institutes of Health; Wellcome Trust; Global Fund to Fight against AIDS, Tuberculosis and Malaria; The Bill and Melinda Gates Foundation.

**Research in context:** *Evidence before this study:* This study describes the genomic analysis of malaria parasites collected during a malaria elimination program targeting four townships (Myawaddy, Kawkareik, Hlaingbwe, and Hpapun) in Kayin State, eastern Myanmar. We searched PubMed on May 10, 2025, using the terms: “malaria”, “transmission”, “drug resistance”, “southeast Asia” and “genetic surveillance” without any date or language restrictions. Our search yielded 58 publications, including reports on declined immunity to malaria and increased drug resistance in southeast Asia; genetic diversity and population structure shaped by mass drug administration, transmission intensity, human movement and mosquito ecology; and the spread of *kelch13* and *pfcrt* mutations conferring dihydroartemimsinin-piperaquine resistance in east southeast Asia countries.

*Added value of this study:* We sequenced and analyzed 2270 *Plasmodium falciparum* whole genomes (each with geographic location) collected between November 2015 and August 2020 in Kayin State. We described fine-scale molecular epidemiology changes during intense malaria elimination interventions. We observed localized, temporally stable transmission of parasite genotypes, with different parasite families and *kelch13* haplotypes predominating in different subregions over the 5-year period. We found *kelch13* haplotypes with local or regional origin, but none originating from eastern southeast Asia. The genomic data indicated decreasing parasite population size, but we observed no selection towards drug-resistance parasites. In 2020, only one predominant lineage (*kelch13*-R561H) remained in our studying region, consistent with genetic drift in a pre-elimination setting.

*Implications of all the available evidence:* The parasite population was genetically depauperate, with a spatially localized and temporally stable distribution of parasite lineages, and declining population size. In this situation genetic drift may play a heightened role in parasite epidemiology. Consistent with this, parasites carrying *kelch13*-R561H rose to high frequency in 2020, because transmission was eliminated in all regions except the north of Kayin State, where parasites bearing this *kelch13* genotype have predominated since 2017. Future surveillance will determine whether this parasite genotype is transmitted to neighboring regions, and thus related to changes in *P. falciparum* drug resistance dynamics on the Thailand Myanmar border. These data illustrate how genetic drift in small populations can result in instantaneous changes in the resistance status of parasite populations in near elimination settings.

## Introduction

Regions of low malaria transmission intensity predominate in Southeast (SE) Asia and South America and are becoming increasingly common in Africa ^1^. A central challenge for malaria control is to develop efficient approaches to eliminate malaria from such regions. Rapid selection of drug-resistant parasites is a central concern for intensive malaria control programs. For example, *kelch13* mutations conferring artemisinin resistance increased from 0-90% frequency in parasites collected from patients visiting Shoklo Malaria Research Unit (SMRU) clinics on the Thailand Myanmar border between 2003 – 2014 ^2^. Use of mass drug administration (MDA) is controversial for malaria treatment due to concerns about resistance. Prior use of chloroquine treated salt is thought to have accelerated selection of chloroquine resistance in the last century ^3^. However, MDA is effective for treating submicroscopic malaria infections which comprise the majority of infections in many SE Asian locations^4^, and are missed by passive malaria surveillance. Submicroscopic infections may be cured effectively by MDA because there are few parasites per patient, and so treatment is more likely to be completely successful than in high-parasitemia infections. It has therefore been argued that MDA is not likely to promote resistance spread in low prevalence regions like SE Asia ^5^. A previous paper examined the epidemiology of *kelch13* haplotypes in Kayin State and showed limited changes of *kelch13* artemisinin resistance alleles between 2013-2019 ^6^. Spread of resistance was much less rapid than occurred in the SMRU clinics (2000-2014); this is consistent with combined use of near-exhaustive coverage of communities with malaria posts and MDA imposing limited selection for ART-resistance.

Here, we examine genomic epidemiology of *P. falciparum* samples collected between November 2015 and August 2020 during malaria elimination efforts in Kayin State. Between 2014-2020, the Malaria Elimination Task Force (METF) targeted four townships (Myawaddy, Kawkareik, Hlaingbwe, and Hpapun) in Kayin State, eastern Myanmar, for malaria elimination ^7,8^. This was done using a combination of interventions: (i) 1475 village malaria posts (MPs) were opened, providing rapid diagnosis and malaria treatment (artemether–lumefantrine plus single low-dose primaquine); (ii) Mass Drug administration (MDA) (dihydroartemisinin–piperaquine [DHA-piperaquine] plus single dose primaquine once per month for 3 consecutive months) was used for 69 “hotspot” villages where malaria remained prevalent (>40% malaria and >20% *Plasmodium falciparum*) ^8^. The combination of these two approaches reduced *P. falciparum* cases by 97% from an incidence of 39 cases per 1000 person-years (May 2014–April 2015) ^8^ to 1 case per 1000 person-years (May 2019 to April 2020) ^9^.

Our central goal was to use genome sequence data to understand parasite transmission, population genomics and resistance evolution, and to use these data to inform future control efforts in elimination settings of SE Asia. These analyses can also help guide future efforts towards malaria elimination in African regions. For example, malaria has been reduced or declared malaria free in some areas of Africa (e.g. Senegal ^1^ and Cabo Verde ^10^), although emerging *kelch13* artemisinin-resistance mutation threatens to derail control and elimination efforts ^11,12^. Our key questions were to: (i) determine the distribution and stability of malaria populations within the Kayin State target region; (ii) to evaluate evidence for long-distance gene flow between Kayin State, Myanmar and other eastern SE Asian countries (Cambodia, Vietnam, Laos); (iii) document the origins of *kelch13* resistance alleles; (iv) to determine the impact of MDA on parasite population structure; and (v) to evaluate appropriate genomic metrics for assessing transmission intensity.

## Methods

### Study area and sample origins

The samples for this analysis were collected during routine diagnosis and treatment efforts in Kayin State as part of the METF malaria elimination effort led by the Shoklo Malaria Research Unit (SMRU, based on the Thailand-Myanmar border) (Figure 1, Appendix Figure 1&2). This METF project was established in 2014 and utilized two primary *P. falciparum*-focused interventions: the establishment of a large network of community-based malaria diagnosis and treatment posts (MPs), and targeted MDA in communities determined to have a high prevalence of asymptomatic *P. falciparum* infections. The MPs were stocked with filter papers (Whatman 3mm blotting paper) and were asked to collect dried blood spots (DBSs) from finger prick blood samples for patients with rapid diagnostic test (RDT) confirmed *P. falciparum* infection. Each DBS sample is linked to the MP from which it originated, and all MPs have geographic references (latitude and longitude). 5014 DBS samples were collected between November 2015 and August 2020 (Table S1). The DBS samples were then transported to SMRU and subsequently shipped to the Texas Biomedical Research Institute (in the U.S.A.) for molecular analyses.

**Figure 1.**
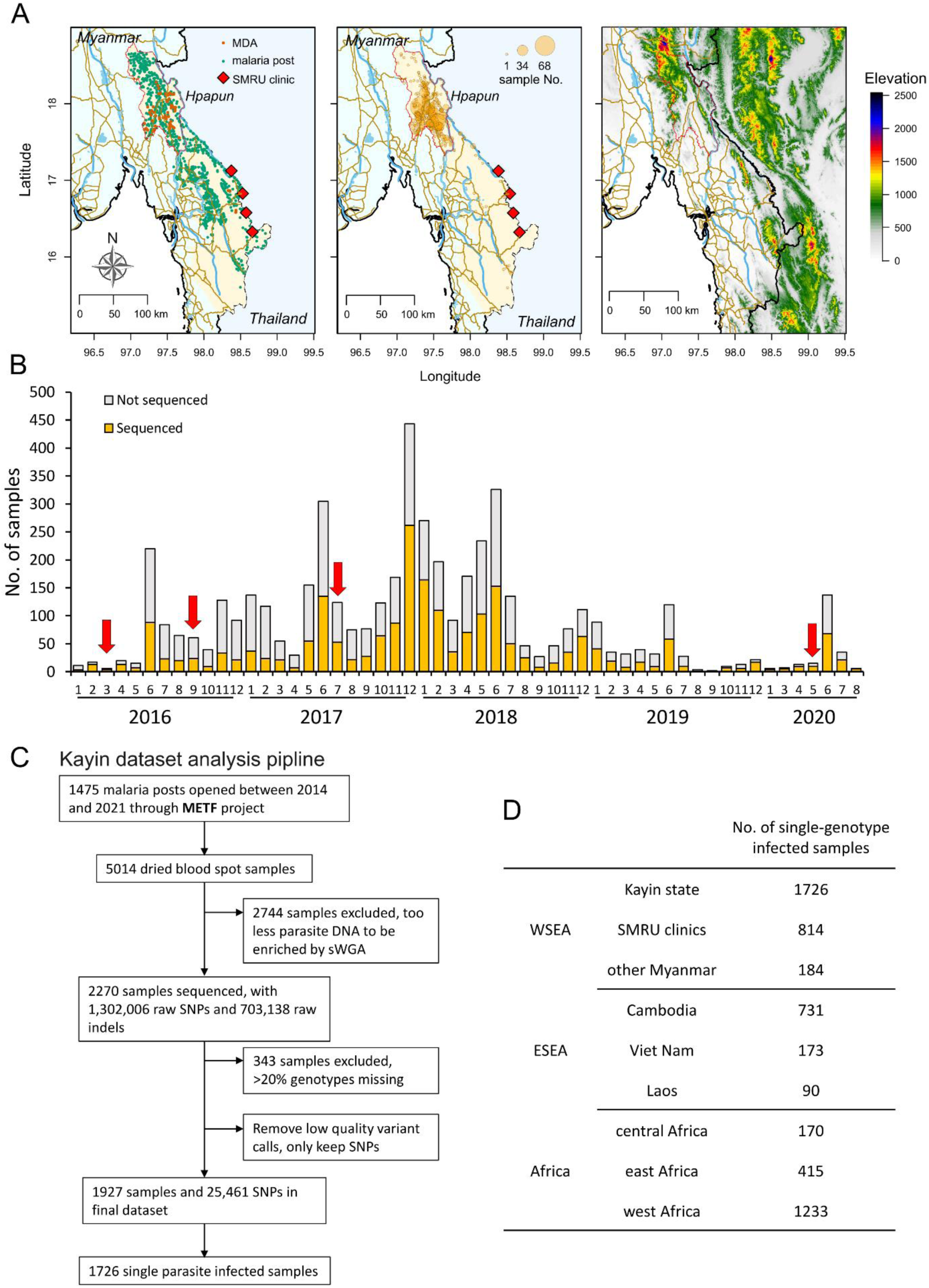
Sample collection and dataset summary. (A) Physical geography of the Malaria Elimination Task Force (METF) intervention region. The METF project was performed at four townships (Myawaddy, Kawkareik, Hlaingbwe, and Hpapun) of Kayin State (light yellow shaded region), Myanmar. Left panel, the distribution of malaria posts (MPs, green dots). Vermillion dots indicate locations where mass drug administration (MDAs) were applied. Middle panel, location of sequenced samples. Over 96% of the sequenced sample were from Hpapun township, the northern part of Kayin State. Right panel, elevation map. Elevation data was downloaded from the United States Geological Survey (USGS, https://earthexplorer.usgs.gov/). The locations of roads (brown) and rivers (light blue) were from Myanmar Information Management Unit (https://themimu.info/). (B) Temporal distribution of samples collected through the METF project. Red arrows indicate the time when MDAs were applied. (C) Analysis pipeline for samples collected from Kayin State. (D) World-wide malaria parasite datasets used in this study. Sequencing data other than those from Kayin State were from MalariaGEN (https://www.malariagen.net/, release 6). Shoklo Malaria Research Unit (SMRU) clinics are located at the Thailand Myanmar border. Samples from Myanmar but not from Kayin State are labeled as “other Myanmar”, see Figure S1 for detailed sample regions.

### Whole genome sequencing and genotyping

We extracted DNA from the dried blood spots and enriched parasite genomes using selective whole genome amplification (sWGA) to following Li et al ^13^ and Oyola et al ^14^ (see **Appendix Methods** for details). sWGA products with more than 50% DNA from *P. falciparum* were used for further library preparation and whole genome sequencing (WGS). We sequenced 2270 DBS samples to an average coverage of 60× using Illumina Hiseq X or Novaseq sequencers. Over 96% of the sequenced samples were from Hpapun Township, in the northern part of Kayin State (Figure 1A, middle panel).

We mapped sequencing reads from each sample against the *P. falciparum* 3D7 genome (PlasmoDB, release 46) and the human GRCh38 genome. Reads mapping to the human genome were discarded before genotyping. We initially identified 1,302,006 single-nucleotide polymorphisms (SNPs) and 703,138 indels (Figure 1C). We removed 343 samples with > 20% genotypes missing. We then filtered the SNP calls following a “stringent” filtering method ^15^, to generate a final list of 447,435 high-quality, biallelic, core-genome located (defined in ^16^) SNPs. To analyze complexity of infection and population structure, we further removed SNPs that were genotyped in less than 50% of samples or with minor allele frequency (MAF) < 0·05.

### Complexity of infection

We measured multiplicity of *P. falciparum* infections using the within-infection *F_ws_* fixation index ^17^. Samples with *F_WS_* > 0·9 were assumed to come from single-genotype infections for samples from Kayin State. Allele frequencies across the genome were plotted and manually inspected to detect further possible complex infections.

### Relationships among parasite genotypes

We used relatedness - *r*, defined as the fraction of the genome that is identical-by-descent (IBD) between a pair of individuals ^18,19^ - to estimate parasite relationships. Based on the distribution of relatedness among F1 progeny from malaria parasite genetic crosses (Appendix Figure 3), we assume that parasites are genetically related if ≥ 25% of their genome is identical (*r* ≥ 0.25); parasites are closely related (such as siblings or parent and progeny) if their relatedness is greater than 45% (*r* ≥ 0.45). We considered samples to be clonal if their relatedness is over 90% (*r* ≥ 0.90). We visualized relatedness among samples using the R package *pheatmap* and the *Cytoscape* software. We also examined the recombination patterns between closely related parasites and plotted shared IBD regions between estimated parents and progeny using *karyoploteR*.

### Surveillance of *kelch13* haplotypes

We extracted SNPs distributed within 100 kb upstream and 100 kb downstream of the *kelch13* gene. We measured expected heterozygosity (*He*) at the *kelch13* locus by treating *kelch13* as a single locus with multiple alleles. We also measured *He* over the 200kb *kelch13* haplotype region. To compare the relationships between different *kelch13* haplotypes, we measured pairwise IBD sharing among all *kelch13* haplotypes. We assume that haplotypes with IBD sharing ≥ 0·90 originated from the same mutational event; that when 0·35 ≤ IBD < 0·90, there was a one least recombination event to break the original haplotype; and when IBD < 0·35, these haplotypes have emerged independently.

### Comparisons of malaria parasite populations

We compared the Kayin State parasite population with other world-wide malaria parasite populations (Figure 1, Table S2). The SMRU clinics are located around Mae Sot, in Tak Province along the international Thailand-Myanmar border. We used “other Myanmar” to represent sampling sites in Myanmar but not from Kayin State. West SE Asia population includes samples from Kayin State, SMRU clinics and other Myanmar regions, while east SE Asia population includes Cambodia, Viet Nam, and Laos.

We merged raw SNP genotypes from the Kayin dataset with those from MalariaGEN *P. falciparum* Community Project ^20^ (release 6). We performed “stringent” filtration as described above, and selected loci with minor allele frequency > 0·05. We calculated genetic richness [R_G_ = (G-1)/(S-1)] ^21,22^ to quantify the richness of clonal parasites in each population, where G is the number of unique genomes, and S is the total number of single genotype infected samples. For samples with relatedness > 0·9, only one representative sample per population with the highest genotype rate was selected and used for further analysis (Table S2). We pruned SNPs for linkage disequilibrium (LD) and generated a pairwise genetic distance matrix using PLINK with default parameters. We conducted principal component analyses (PCA) and ADMIXTURE analyses based on the pruned genotypes and distance matrix. We measured the proportion of pairs IBD across the genome within populations following the scripts in *isoRelate* ^23^. We estimated effective population size (*Ne*) based on patterns of LD at unlinked loci, using methods implemented in *NeEstimator* v2·0 ^24^.

### Statistical analysis

All statistical analysis was performed using R version 4·1·3. For pairwise comparisons between groups, we used Welch Two Sample T-test. We measured correlations between parasite genetic relatedness and geographic distance or time using the Mantel statistic using the *mantel* function in the *vegan* package. *p*<0·05 was considered statistically significant. We used hierarchical clustering on principal components (HCPC) following scripts in *FactoMineR* ^25^ to divide the 283 malaria posts with samples sequenced into 50 HCPC regions based on latitude and longitude. We then compared parasite relatedness within and between HCPC regions for parasites collected in the same year, between 1-2, 2-3 and 3-4 years apart. We compared relatedness between parasites collected from HCPC regions 6 months before and 6 months after MDA. As controls, we examined relatedness of parasites collected from HCPC regions where MDA was not used during the same time windows.

## Results

### Extreme clonal expansion and inbreeding in a region under massive drug selection

We processed and analyzed genome-wide sequencing data of 2270 DBS samples collected from 283 malaria posts in eastern Myanmar, between November 2015 and August 2020 (Table S1, Figure 1). After filtration of low-coverage samples and low-quality genotypes, the final Kayin State dataset contains 1927 *P. falciparum* samples with a set of 25,461 high-quality SNPs. 89·6% (1726/1927) of the samples were from single-genotype infections (*F_ws_* ≥0·90, Appendix Figure 4).

**Figure 2.**
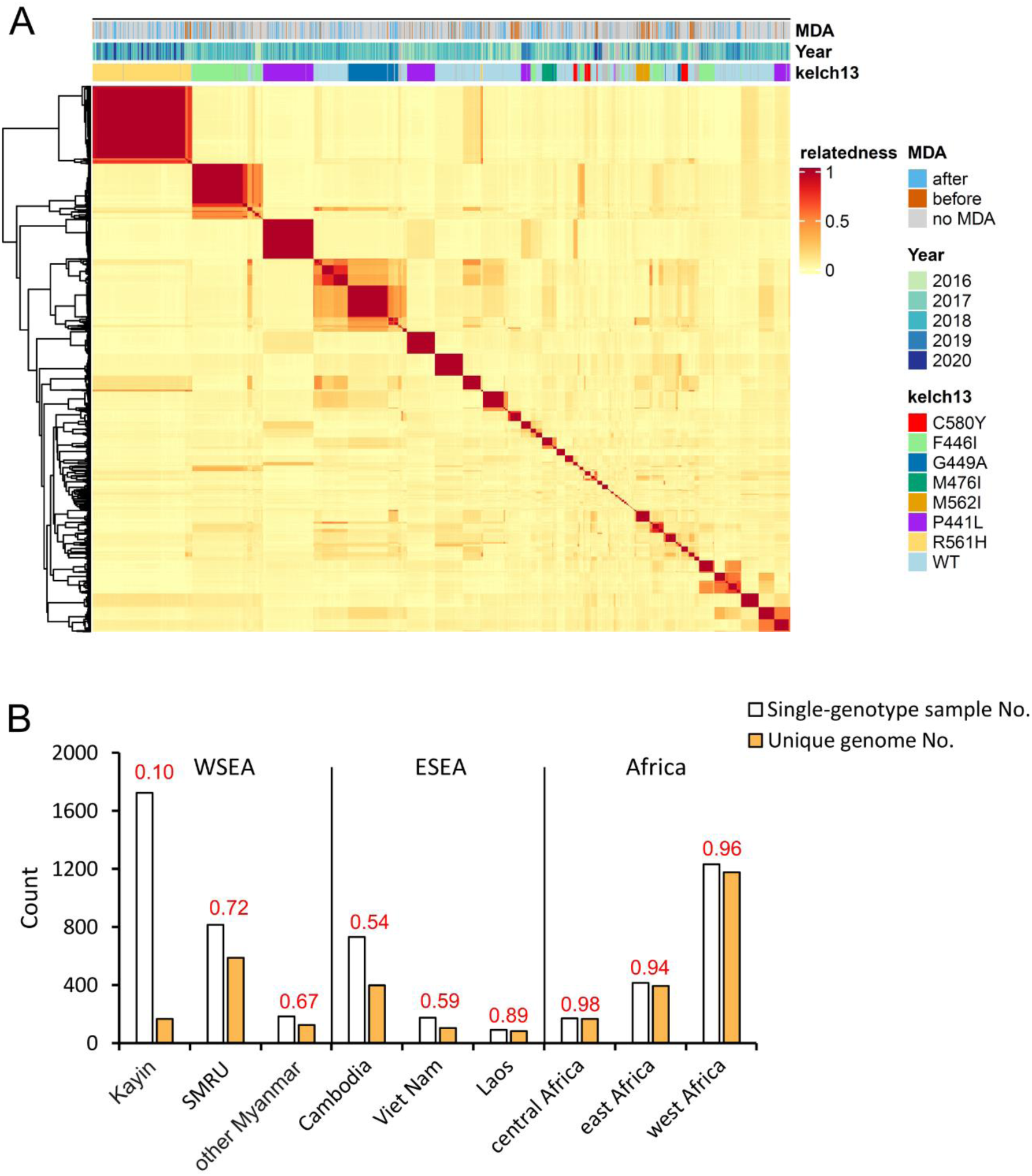
Parasite relatedness and level of clonal transmission. (A) Heatmap showing relatedness among Kayin samples. Pairwise parasite relatedness (*r*) was measured as the proportion of genomes that are identical by decent (IBD) between pairs of samples. Samples with *r* ≥ 0·9 are considered as IBD and share the same unique genome. Color bars at the top of the heatmap indicate information for each sample: MDA, if the sample came from a malaria post with (orange) or without (blue) mass drug administration; Year, the year of sampling; *kelch13*, the genotype of *kelch13* - only alleles with frequency > 2% in at least one sampling year were colored. (B) Level of clonal transmission. Red numbers on top of bars indicate genetic richness. Kayin population had the highest level of clonal transmission compare to other populations. SMRU, Shoklo Malaria Research Unit; WSEA, west southeast Asia; ESEA, east southeast Asia.

**Figure 3.**
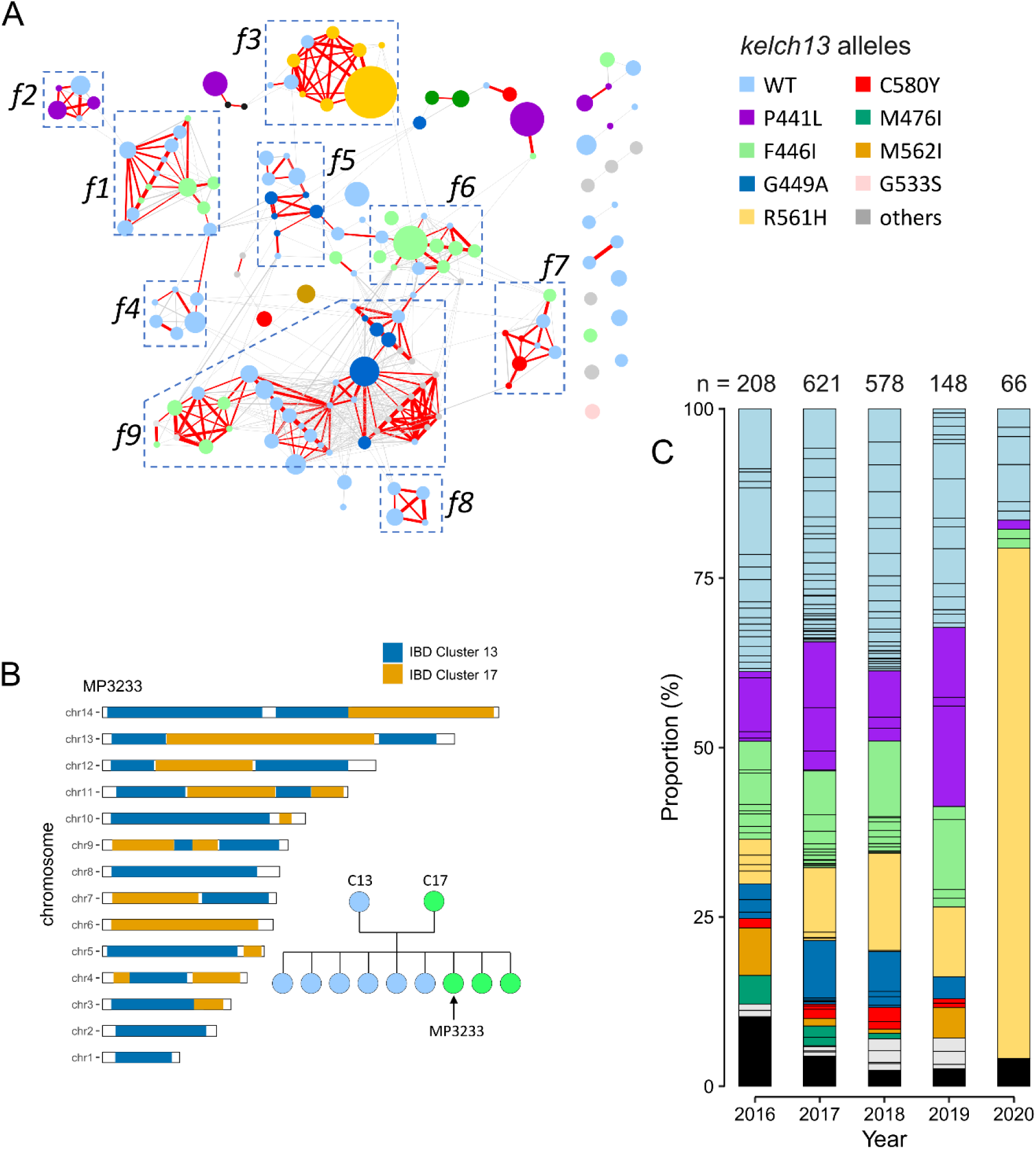
Parasite clonal expansion and inbreeding in Kayin State. (A) IBD network of unique genomes from Kayin population. Nodes: each circle indicates one unique genome and is color coded based on its *kelch13* alleles; circle size indicates sample size (ranged from 1 to 229). Edges: connections with relatedness (*r*) ≥ 0·25; thicker lines indicate higher relatedness; red lines are connections with *r* ≥ 0·45. Parasites from closely related families (f1 to f9) are labeled using boxes. 152 of 166 unique genomes were include in the network, representing 98·6% of single-genotype infected samples. (B) Pedigree tree of parasites from family 1 (f1) and chromosome plot for an estimated progeny (MP3233). See Figure S7 for chromosome plots for all progeny. We infer that the parents of f1 are C13 (IBD cluster 13, *kelch13-*wildtype) and C17 (IBD cluster 17, *kelch13*-F446I). (C) Proportion of unique genomes across time. Each segment within a bar represents one unique genome, which is colored based on its *kelch13* allele. Black blocks indicate number of unique genomes that were recovered only once (“singletons”). A clonal expansion of IBD cluster 1(*kelch13*-R561H) parasites was detected in 2020.

**Figure 4.**
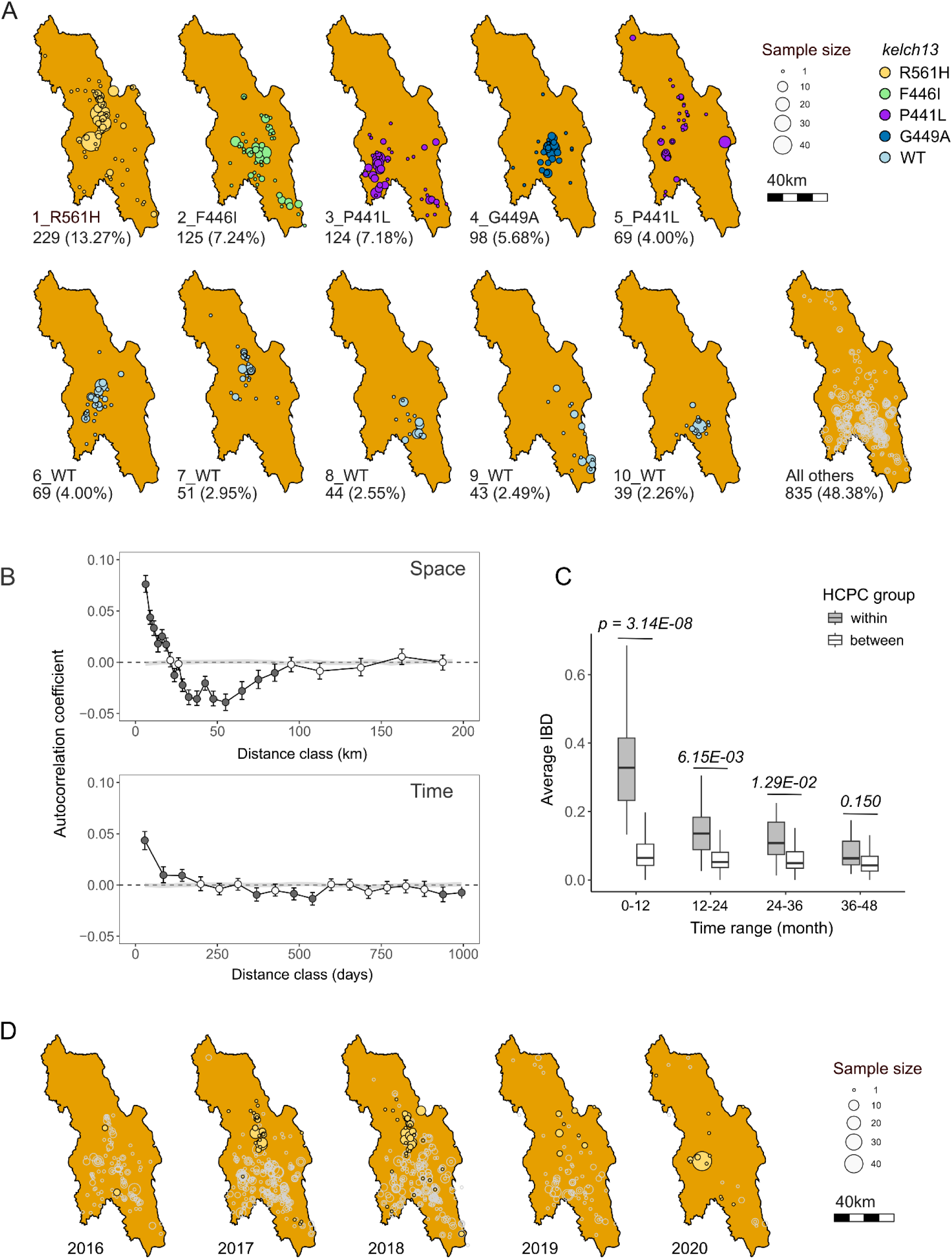
Localized transmission and temporal stability. (A) Spatial distribution of different IBD clusters. Each circle represents samples collected from an individual malaria post. IBD cluster IDs and corresponding *kelch13* alleles are indicated at the bottom left of each panel; for example, 1_R561H indicates IBD cluster number 1 carrying the *kelch13* R561H allele. (B) Correlogram analysis of pair-wise parasite relatedness across space and time. (C) Comparison of relatedness between within-group and between-group HCPC region pairs. X-axis indicates time intervals in month. (D) Temporal dynamics and spatial distribution of the IBD cluster 1 (*kelch13* R561H) through the sampling year.

To identify highly related individuals, we estimated pair-wise genetic relatedness (*r*, proportion of genome that was IBD) and used these to cluster samples with ≥90% of the genome IBD (*r* ≥0·90) (Figure 2). From the 1726 single-genotype Kayin samples, we identified 93 IBD clusters with unique genomes (2 to 229 samples per cluster), and 73 singletons, giving a total of 166 unique genomes. The Kayin population has an R_G_ of 0·10 (166 unique genomes from 1726 single-genotype infected samples), containing 14 large clonal expansion clusters (>30 samples per cluster, Figure 2). In contrast, the R_G_ ratios are much higher for the other SE Asian populations, with 0·72 (589/814) for SMRU clinics, 0·67 (123/184) for other Myanmar regions (not include Kayin), 0·54 (398/731) for Cambodia, 0·59 (102/173) for Viet Nam and 0·89 (80/90) for Laos. R_G_ ratios range from 0·94 to 0·98 for African parasite populations (Figure 2B).

Samples from the most common 10 IBD clusters account for 51·62% of the population, showing low parasite diversity in Kayin State. 151 out of 166 (91·0%) unique genomes were genetically related (*r* ≥0·25) to at least one other unique genome, indicating high levels of inbreeding in the Kayin State parasite population (Figure 3A). We identified 9 closely related families, that account for 110/166 unique genomes. Each individual in a family has a *r* ≥0·45 with at least another family member. Samples from these families account for 65·2% (1126/1726) of all the single-genotype infections in Kayin State. To further estimate genealogical relationships of individuals inside each family, we analyzed the distribution of chromosomal IBD segments between closely related family members (Figure 3B, Appendix Figure 5). For two of the families (family 1 and family 7), we were able to identify both parents and F1 progeny based on their chromosomal recombination patterns, indicating extremely small parasite population size in Kayin State.

**Figure 5.**
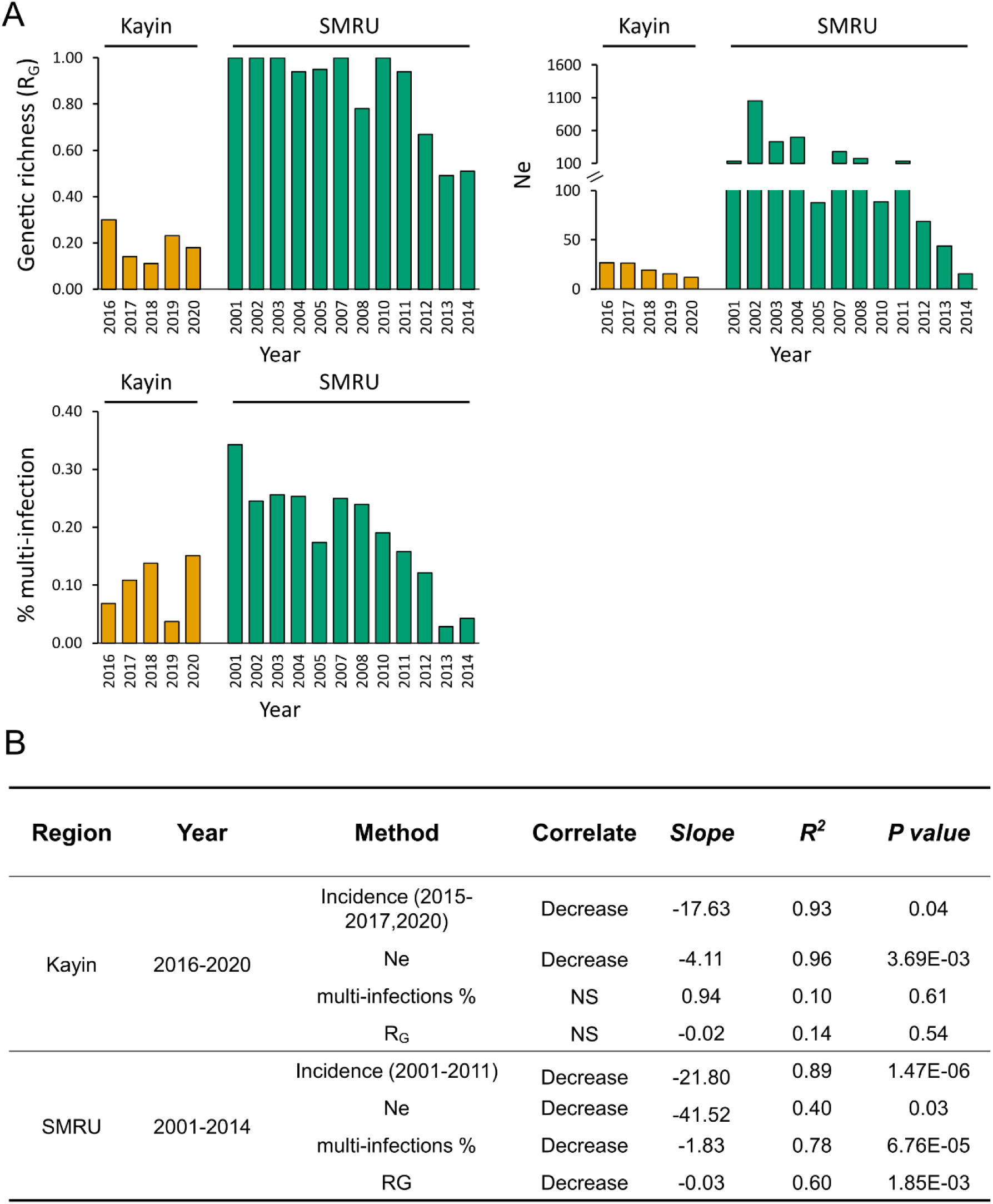
Genomic measures of parasite population size. (A) Population size estimated using genetic richness (RG), proportion of samples from multiple-genotype infection (multi-infections%), and effective population size (Ne). (B) Comparison of genetic metrics between Kayin State and Thailand-Myanmar border (SMRU clinics) populations. The incidence data for Kayin State was from Landier et al., 2018 and Legendre et al., 2023; and the incidence data for SMRU clinics (regions near Mae Sot) was from Nkhoma et al., 2013. NS, not significant; SMRU, Shoklo Malaria Research Unit clinics.

### Localized transmission and regional stability of haplotypes

#### Genetic relatedness in space

Spatial groups of IBD clusters reflect direct or indirect transmission chains of malaria parasite clones, so are particularly informative for understanding transmission dynamics. The IBD clusters and closely related families show localized spatial distribution (Figure 4A, Appendix Figure 6). For example, 86% samples from the largest IBD cluster (carrying *kelch13* R561H) were collected to the north of Hpapun Township. The second largest IBD cluster (carrying *kelch13* F446I) was found mainly in the center of Hpapun Township, while the third largest IBD cluster (carrying *kelch13* P441L) was found in the west of the same township. Similarly, different IBD clusters carrying *kelch13* wildtype alleles, show localized distribution (Figure 4A). Spatial correlograms confirm that parasite relatedness is positively correlated at distances ≤20 km (Figure 4B). We also observed significant negative correlations in relatedness between 27·5-90km. The negative correlations further indicate the spatial pattern of parasite relatedness. Not only are parasites that are geographically proximate more likely to be related – those that are geographically distal (27·5-90km) are less likely to be related.

**Figure 6.**
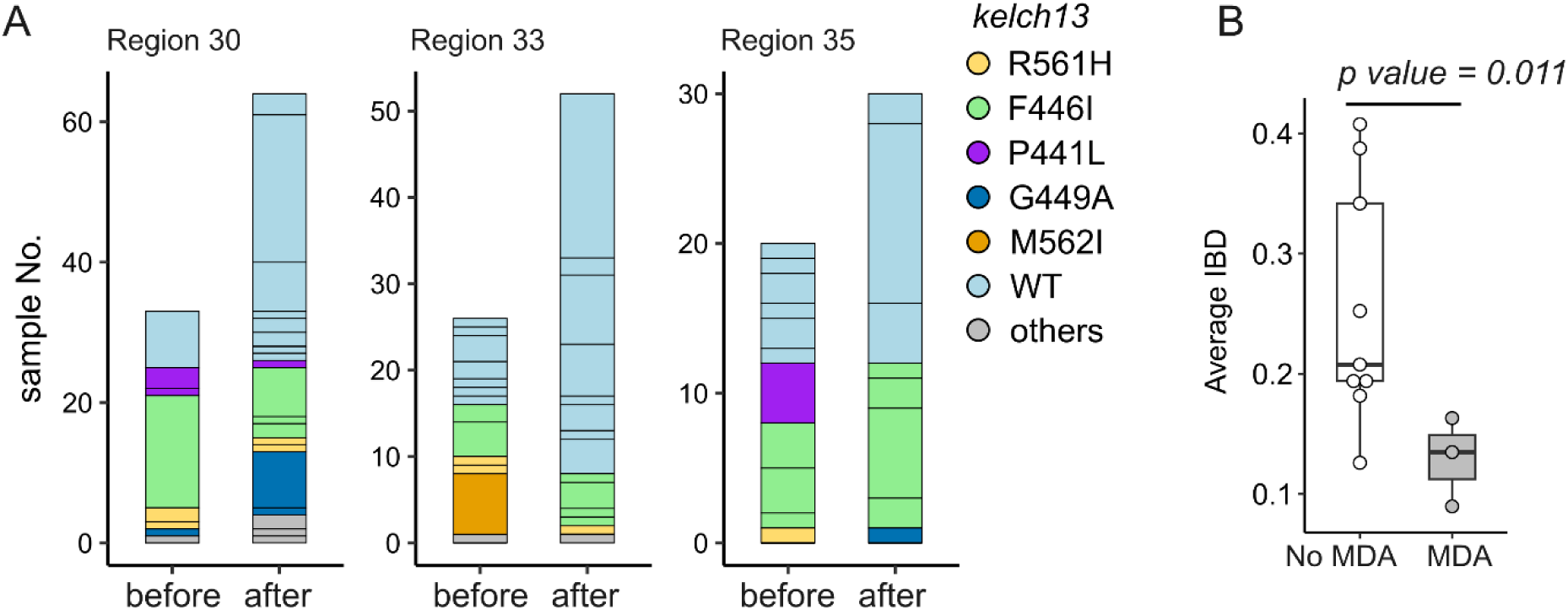
The impact of mass drug administration (MDA). (A) Population relatedness before and after MDA. MDAs were implemented between June 24, 2017 – October 12, 2017 in HCPC regions 30, 33 and 35, using DHA-piperaquine plus a single dose of primaquine administered monthly for three consecutive months. Bar segments represent unique genomes identified within the 6 months before (April 12, 2017 – October 12, 2017) or after the intervention (October 13, 2017 – April 12, 2018), color-coded by *kelch13* allele type. (B) Comparison of parasite genetic relatedness between HCPC regions with and without MDA intervention. Parasites collected from the same time windows from HCPC regions where MDA was not used provided “no MDA” controls.

#### Genetic relatedness in time

The length of time in which clonal lineages are sampled provides an indication of the frequency of outbreeding within malaria parasite populations. We detected clonal IBD clusters (*r* ≥ 0·90, contain ≥2 samples) that were sampled across the 56-month study period, as well as new genome haplotypes generated through recombination (Appendix Figure 7). Out of the 93 IBD clusters, 9 lasted ≥ 36 months (3 years of sampling), while 34 lasted ≤ 6 months. The mean duration of IBD clusters in Kayin population was 13·8 (1 *se* = 1·4) months. Furthermore, the mean sampling duration of closely related parasite family members (*r* ≥0.45) was 48·7 (1 *se* = 3·0) months (Appendix Figure 7), consistent with a low frequency of outbreeding in the population. We further analyzed the two parasite families for which both parents and progeny were identified. There were only 12 recombination events among the 109 samples (14 unique genomes, 41 months of family duration) from family 1, and 6 among the 31 samples (8 unique genomes, 48 months of family duration) from family 7 (Table S1, Appendix Figure 5), consistent with low parasite outbreeding frequency.

We used a temporal correlogram to examine the number of days over which correlations in relatedness were observed (Fig 4B). This revealed positive correlations in relatedness between parasites sampled ≤ 170 days apart. To evaluate the relative impact of space and time on parasite relatedness, we divided the control region into 50 regions using hierarchical clustering on principal components (HCPC) (Appendix Figure 8); 29 of these HCPC regions contained from 10-161 parasites. We examined relatedness in parasites sampled within and between HCPC regions for parasites sampled parasites sampled 1-12, 13-24, 25-36 and 37-48 months apart (Figure 4C). We observed significantly greater relatedness among parasites sampled from the same HCPC unit, relative to those sampled from different HCPC units. This remained significant for parasites sampled up to ≤36 months apart, demonstrating spatial stability of parasite populations within Hpapun Township (Figure 4C).

### Long distance connectivity of parasite populations in SE Asia

We used parasite relatedness to measure connectivity within west SE Asian populations and between west and east SE Asia (Appendix Figure 9). We detected a high level of gene flow between parasite populations from SMRU clinics and Kayin State: 38·3% of Kayin samples had >25% IBD (*r* > 0·25) with at least one sample from SMRU clinics, and 18·1% had >35% genome IBD (*r* > 0·35). We identified two subpopulations, corresponding to west SE Asia (Kayin State, SMRU clinics and other Myanmar regions) and east SE Asia (Cambodia, Viet Nam and Laos) based on their genetic similarity and population structure (Appendix Figure 10). Hence, we found no evidence for clonal transmission or recent recombination (*r* > 0·15) between west and east SE Asia.

The low connectivity between west and east SE Asia is further confirmed by the distribution of *pfcrt* mutations conferring piperaquine (PPQ) resistance. 52·19% of east SE Asian (Cambodia, Viet Nam and Laos) parasites carried PPQ-resistant *pfcrt* alleles between 2015 and 2018 (MalariaGEN Pf7 ^26^). These included T93S (23·55%), I218F (11·87%)^27,28^, H97Y (5·37%), F145I (8·20%), G353V (2·125)^29^ and G367C (1·08%)^30^. In contrast, these *pfcrt* mutations were absent from the Kayin dataset (Table S1) and from other west SE Asia regions (other Myanmar regions and west Thailand).

### Genomic measures of parasite population size

We evaluated three genetic metrics (proportion of multiple-genotype infections, R_G_, and *Ne*) in both Kayin State and SMRU parasite populations, for assessing how control efforts impact parasite population size (Figure 5). The malaria incidence decreased significantly in both regions over studying time. The incidence decreased from 273·9 cases per person-year in 2001 to 22·4 in 2011 for SMRU clinics ^31^, and from 58·8 in 2016 to 1·0 in 2020 for Hpapun Township in the northern part of Kayin State, where >96% sequenced samples were collected ^8,9^ (Table S3).

#### Proportion of multi-infections

The proportion of multiple clone infections (10·4%, 201/1927) in Kayin samples was low compared to other *P. falciparum* populations (Table S3) and remained low throughout the year (range: 3·7-15·1%). The proportion of multiple clone infections decreased significantly in SMRU clinics, from 34·3% in 2001 to 4·3% in 2014 (*p*-value = 6·76e-05, *R*^2^ = 0·78), consistent with a prior analysis ^31^. However, this statistic showed no significant decline in Kayin (*p*-value = 0·61, *R*^2^ = 0·10, Figure 5B).

#### R_G_

R_G_ ratio is expected to be negatively correlated with the level of clonal expansion and positively correlated with transmission intensity ^31^. R_G_ ratios were lower in Kayin (0·10 to 0·30) between 2016 and 2020 than in SMRU clinics (range: 0·50-1·00) between 2001-2014. The R_G_ ratio decreased over time in SMRU clinics (*p*-value = 1·85e-03, *R*^2^ = 0·78), but not in Kayin (*p*-value = 0·54, *R*^2^ = 0·14)

#### Ne

We computed single-sample estimates of effective population size (*Ne*) using unique genomes from each population for each sampling year. *Ne* estimates were lower in Kayin (11·5 to 26·6) compared to SMRU clinics (15.5 to infinite). We detected significant reductions in *Ne* in both Kayin (*p*-value = 3·69e-03, *R*^2^ = 0·96) and SMRU clinics (*p*-value = 0·03, *R*^2^ = 0·40).

### The evolution of *pfkelch13* alleles

#### Impact of malaria elimination efforts on drug resistance

61·32% of samples from Kayin State carried nonsynonymous SNP mutations in *kelch13* (Appendix Figure 11). The major mutant alleles were P441L (15·19%), F446I (15·01%), R561H (14·02%), and G449A (7·87%). Only 2·28% of Kayin samples carry C580Y. In comparison, in the adjacent SMRU clinics, C580Y was the dominant *kelch13* mutant allele, reaching 71·05% allele frequency in 2014 (Table S2). There were 47 IBD clusters (*r* ≥ 0·90) carrying mutant *kelch13* alleles and 46 IBD clusters carrying wild-type *kelch13*. We compared the size of IBD clusters carrying mutant *kelch13* alleles with those carrying wild type *kelch13* and found no significant difference (Figure 3C, Appendix Figure 12). These results suggest that artemisinin selection was not the main driver for clonal expansion in Kayin State.

#### Clonal expansion of parasite carrying *kelch13*-R561H in 2020

Despite strong drug selection, frequencies of mutant *kelch13* alleles remained stable between 2016 and 2019 ^6^ (Figure 3C, Appendix Figure 12). However, in 2020 one of the *kelch13* alleles - R561H - reached 74·2%. This clonal expansion results from near elimination of parasites from most areas in Kayin, with the exception of northern Hpapun Township where parasites carrying *kelch13*-R561H predominate (Figure 4D). 54·8% (40/73) of samples collected between January and August 2020 before the COVID-19 pandemic lockdowns were from one single malaria post (LH-0266B) (Table S1).

The change in *kelch13* allele frequencies were reflected by changes in diversity in this gene and its flanking regions. We measured expected heterozygosity (*He*) to quantify diversity. *He* in *kelch13* remained high between 2016-2019 (*He* = 0·78) but dropped to 0·41 in 2020. We observed parallel reductions in flanking region diversity with a drop from 0·48 to 0·19 between 2019 and 2020 (Appendix Figure 11D).

#### Origins of *kelch13* alleles

We reconstructed the haplotypes surrounding the *kelch13* gene (100kb upstream and 100 kb downstream). We found a wide variety of *kelch13* genetic backgrounds, with one or more unique haplotypes per resistance allele (Appendix Figure 13 & 14). Two P441L, one F446I, one G449A and one R561H haplotypes had shared ancestry between Kayin State and SMRU clinics or other Myanmar regions. However, none of these alleles had high frequency in SMRU clinics or other Myanmar regions. Two F446I and one C580Y haplotypes were uniquely observed in Kayin State, consistent with local origin. For two *Kelch13* resistance alleles (G449A, R561H), the wildtype *kelch13* haplotypes on which these resistance mutations arose were sampled in the early 2000s in SMRU clinics. Of the three C580Y haplotypes identified in Kayin, two were also found in SMRU clinics, while one was unique to Kayin State. The two C580Y haplotypes shared with SMRU clinics were only found to the south of Hpapun Township and 60-120km north of the SMRU clinics (Appendix Figure 15). None of these haplotypes shared IBD with east SE Asia C580Y haplotypes (Appendix Figure 16).

### Impact of mass drug usage on parasite population structure

We predicted that MDA would reduce relatedness of pre and post MDA parasite populations due to clearance of the local parasite population and replacement with new parasite genotypes post-MDA, resulting in founder effects. There were three HCPC units in which sufficient parasites (n ≥ 20) were sequenced both pre and post MDA (Figure 6A). The relatedness between pre and post MDA parasites from these 3 HCPC units was significantly lower than observed between malaria parasites collected during the sampling time period from HCPC regions where MDA was not used (Figure 6B). Hence, MDA impacted parasite relatedness, consistent with efficient control of pre-MDA parasite genotypes, and post-MDA recolonization with unrelated genotypes.

## Discussion

The METF elimination efforts, combining community malaria posts and MDA, significantly decreased malaria case numbers in the target area (near the Myanmar-Thailand border) between 2014-2020 ^8,9^. We described the key results from genomic surveillance during these elimination efforts, including parasite transmission patterns, population diversity and genomics, evolution of drug resistance, and genomic impacts of MDA using over 2000 whole genome sequenced *P. falciparum* samples collected between Nov 2015 – Aug 2020.

### Spatial and temporal structure of malaria populations

Parasite sequences from the METF study region reveal extremely high levels of inbreeding and low levels of genetic variation. We found 166 genotype clusters (≥90% of the genome IBD) among the 1726 single clone samples sequenced. Hence only 10% of parasite genomes sampled are unique (R_G_ = 0·1), and most infections show a clonal structure. In contrast, genetic richness is >0·94 in African populations sampled, and ranges from 0·54 to 0·89 in other SE Asian populations examined ^20^. Furthermore, 110 of the 166 unique genomes are distributed among 9 different families (r ≥ 0·45). Parasites within these families typically carry one or two *kelch13* alleles, that mostly likely inherited from the parents. This is clearly the case in the two families with both parents identified - family 1 (F446I and wild-type *kelch13*) and family 7 (C580Y and wild-type *kelch13*). Hence recombination is rare in these populations and most infections are clonally related.

We observed strong spatial structure in the parasite population. This is particularly clear from the distribution of unique parasite genotypes. Such parasites result from self-fertilization of male and female gametes of the same genotype and allow spatial tracing of transmission networks. That these transmission networks are clustered in space is clear visually (Figure 4A), and statistically evident from autocorrelation analyses (Figure 4B), which reveal positive correlations in relatedness between genotypes for up to 20km. The local distribution of unique genotypes indicates (i) local transmission, and very few long-distance transmission events and (ii) reintroduction of circulating genotypes from asymptomatic carriers.

Unique parasite genotypes were long lived in this low transmission setting, with some IBD clusters observed over the complete 56-month study period. This is clear from (i) the temporal autocorrelation, which reveals positive correlation in relatedness between parasites collected 170 days apart; (ii) from the elevated relatedness observed in parasites collected from the same HCPC regions but up 3 years apart. The strong spatial and temporal sub-structure of parasite populations in Kayin State is comparable to that observed in Cambodia ^32^ and Guyana ^33^.

### A clonal expansion of parasites carrying *kelch13*-R561H in 2020

As elimination approaches, genetic drift is expected to play an increasingly important role and expansions of parasite lineages may occur ^34^. We observed a sudden increase in the frequency of IBD cluster 1 parasites carrying *kelch13*-R561H in 2020. In this case, the increase of *kelch13*-R561H frequency resulted from elimination of malaria from all regions of Kayin other than northern Hpapun Township (Figure 4A), where IBD cluster 1 carrying *kelch13*-R561H is at high frequency. Despite predominating in northern Hpapun Township since 2017, the *kelch13*-R561H didn’t spread into other areas of Kayin State. The rapid frequency increase of IBD cluster 1 in 2020 is consistent with bottlenecks and genetic drift in a *P. falciparum* population nearing elimination. Further sampling will determine whether this parasite genotype spreads further in Kayin State and elsewhere in Myanmar and Thailand. In contrast, Wasakul et al ^34^ describe a classic outbreak driven by a selective sweep in Laos, where a lineage carrying *kelch13*-R539H (named LAA1) rose from a low frequency to replace the previously dominant KEL1/PLA1 (C580Y) population.

### MDA impacts parasite population structure

Two control measures were used in Kayin State by METF: malaria posts (early diagnosis and community case management), and regional MDA in malaria hotspots ^8^. The combination of these approaches significantly decreased malaria incidence ^8,9^. Encouragingly, there was minimal evidence of selection for drug-resistance parasites through these elimination efforts which agrees with observations by Imwong et al ^35^, McLean et al ^36^ and Thu et al ^6^. At the genomic level, these combined control measures reduced parasite effective population size (Figure 5).

This study provided an opportunity to examine the impact of one component of this elimination strategy - MDA - on parasite population structure. MDA is expected to generate bottlenecks between pre and post MDA malaria populations, because reservoirs of asymptomatic malaria are removed. We therefore expect to see founder effects resulting from newly colonizing parasites and large divergence between pre and post MDA populations, when compared to populations with no MDA. Our power to detect an impact was limited because most parasites sequenced were collected after MDA: only three HCPC units had sufficient numbers of infection sampled both pre and post MDA to examine this hypothesis. Nevertheless, as predicted, we saw lower relatedness between pre and post treatment parasites in these three HCPC regions than in control regions where MDA was not implemented. These results provide genomic evidence for the effectiveness of MDA in removing local parasite populations through effective clearance of both asymptomatic and symptomatic parasite infections.

### Genomic metrics for assessing transmission intensity

Genetic metrics, such as proportion of multiple clone infections can provide useful metrics for assessing control efforts ^31,37,38^. Such metrics are particularly useful in low transmission regions, where high prevalence of low-density asymptomatic infections complicates assessment of transmission using standard epidemiological methods. However, genetic metrics perform poorly when transmission levels are extremely low. In Senegal ^37,38^, complexity of infection provided an unreliable metric for evaluating transmission when transmission is incidence < 100 cases per 1000 person-years. Consistent with this, we observed that both R_G_ and proportion of Multiple infections worked poorly in Kayin where incidence ranged from 1 - 39 cases per 1000 person years. However, we found that another metric, effective population size (N_e_) calculated from LD between unlinked markers, showed significant decline in both the Kayin State parasite population and from SMRU clinics. We suggest that N_e_ may be a useful genomic indicator of transmission dynamics, particularly in parasite populations in which transmission has been reduced to near elimination levels. N_e_ is typically used in conservation biology to assess viability in endangered populations of animals and plants. Our results suggest that this metric may also have utility for assessing whether parasite populations are approaching local extinction.

### No evidence for long-distance gene flow between Kayin State and Eastern SE Asian countries

The current frontline treatments for *P. falciparum* malaria parasites have been failing in east SE Asia ^35,39,40^, due to the spread of the multidrug-resistance parasites carrying *kelch13*-C580Y mutation and *plasmepsin* 2 amplifications, named KEL1/PLA1. The KEL1/PLA1 lineage was first detected in Cambodia as the DHA-piperaquine was heavily used. When Cambodia withdraw DHA-piperaquine and adopted to artesunate–mefloquine, KEL1/PLA1 subgroups with acquired *pfcrt* mutations conferring piperaquine resistance rapidly spread to other ESEA countries, such as Laos and Vietnam ^39^. There is a concern that this parasite lineage will further spread to west SE Asia, which has the majority of malaria cases in SE Asia ^36^ and where DHA-piperaquine is the frontline treatment for *P. falciparum*. Two lines of evidence suggest minimal geneflow between east and west SE Asia. First, we did not detect *pfcrt* mutations associated with piperaquine resistance on Thailand-Myanmar border or in the Kayin State sampling sites. Second, examination of genome-wide IBD sharing among 3718 infections (1458 unique genomes) revealed no recent recombination or clonal transmission between east and west SE Asia (Appendix Figure 9).

### Origins of kelch13 resistance alleles

*kelch13* mutations conferring artemisinin resistance are established in both east and west SE Asia. C580Y is the major mutation in regions other than northern Myanmar, where F446I predominates ^35^. In contrast, the dominant *kelch13* mutations in the Kayin State include P441L, F446I, R561H, and G449A, depending on the location (Figure 4). While the majority of infections from nearby SMRU clinics carry C580Y (71.05% in 2014), the C580Y frequency in Kayin State was only 2.28%. The most widespread F446I haplotypes in Kayin State originated independently from the dominant F446I haplotype in northern Myanmar.

What factors lead to the patterns of artemisinin resistance evolution seen in Kayin? Longitudinal studies in both Cambodia and from SMRU clinics have revealed that multiple independent *kelch13* mutations emerged and spread initially (soft selective sweeps). Single *kelch13* genotypes (typically *kelch13*-C580Y) eventually outcompete other lineages leading to hard selective sweeps ^2,32,35,39^. In contrast, we found limited evidence that strong drug selection drove drug resistance evolution in Kayin State: (i) we found no significant increase in *kelch13* mutant allele frequencies before 2020 (Figure 3C, Appendix Figure 11) ^6^; (ii) the size of clonal clusters was not significantly different when comparing *kelch13* wildtype and mutant parasites (Figure 3C, Appendix Figure 12). The small effective population size of malaria parasite populations may contribute to the patterns observed, because selection is inefficient when population sizes are small and genetic drift is enhanced ^41^. The initial effective population size of malaria parasites in the Kayin State dataset was much smaller (Ne = 11·5 to 26·6) compared to SMRU clinics (15.5 – infinite) (Figure 2, Figure 5). Other factors that may also limit the impact of drug selection. Human population movements were more limited in Kayin compared to nearby SMRU clinics, especially in Northern Hpapun Township where human movement is limited by difficult terrain, the heavily militarized landscape, and a lack of year-round roads ^7,42^, which can hinder transmission of resistance alleles. Similarly, low levels of recombination in Kayin State limits the rate of formation of new multi-locus parasite genotypes. The small parasite population size, limited population movement, and minimal recombination enhance the role of genetic drift rather than selection in determining drug resistance evolution in the Kayin State region. Our results, and those from other studies ^35^ illustrate how genetic drift can result in rapid stochastic changes in parasite population genomics and drug resistance status in small parasite populations close to elimination.

This study had several limitations: (i) We analyzed malaria genomes collected from 2015 onwards. However, control efforts began earlier than this in 2014. Hence, we were unable to examine malaria population structure and diversity prior to initiation of control efforts. (ii) Use of finger prick blood samples and whole genome amplification resulted in bias towards sequencing high parasitemia infections. (iii) we were unable to score copy number variants, in genes such as Plasmepsin II/III, associated with piperaquine resistance from whole genome amplified DNA. However, the sustained sampling of a high proportion of blood spots collected over a 5-year period provides a unique dataset for examining impact of malaria control efforts on parasite population structure and resistance evolution.

## Contributors

XL, FN, and TJCA conceived and designed the study. JL, AMT, GD, DMP, KML, KS and FN coordinated sample and data collection. XL, GAA, and AR processed samples, and generated genomic data. XL analyzed and interpreted the sequencing data, with input from TJCA, DMP, JL and FN. KML, KS, JL, DMP, and FN were involved in the management and coordination of the genetic surveillance project. XL and TJCA wrote the initial manuscript. XL, DMP, JL, FN and TJCA revised the manuscript. XL, DMP, FN and TJCA accessed and verified all the data. All authors provided critical revision of the manuscript. All authors had full access to all the data in the study and had final responsibility for the decision to submit for publication. TJCA and FN contributed equally.

## Data sharing

Raw sequencing data for the 2270 sequenced samples collected by the Malaria Elimination Task Force project from Myanmar used in the present analysis have been submitted to the NABI Sequence Read Archive (SRA, https://www.ncbi.nlm.nih.gov/sra) under the project number of PRJNA864839. The analysis code and data matrices (genetic distances, geographic distances and temporal distances) are available at: https://github.com/emilyli0325/Malaria-genomics-in-Eastern-Myanmar. This publication also uses data from the MalariaGEN *P falciparum* Community Project ^20^.

## Acknowledgments

This work was supported by National Institutes of Health (NIH) grant R37 AI048071 (to TJCA) and P01 AI127338. Work at Texas Biomedical Research Institute was conducted in facilities constructed with support from Research Facilities Improvement Program grant C06 RR013556 from the National Center for Research Resources. SMRU is part of the Mahidol Oxford University Research Unit supported by the Wellcome Trust of Great Britain. The malaria elimination program (Malaria Elimination Task Force, METF) in Kayin State, Myanmar is supported by the Regional Artemisinin Initiative (Global Fund to Fight AIDS, Tuberculosis and Malaria) and the Bill and Melinda Gates Foundation (OPP1117507). The authors would like to acknowledge the contribution from all member of METF and SMRU, collaborators, and colleagues who have supported the elimination program.

